# Robust Antibacterial Activity of Tungsten Oxide (WO_3-X_) Nanodots

**DOI:** 10.1101/494260

**Authors:** Guangxin Duan, Lu Chen, Zhifeng Jing, Phil De Luna, Lin Wen, Leili Zhang, Lin Zhao, Jiaying Xu, Zhen Li, Zaixing Yang, Ruhong Zhou

## Abstract

Antibacterial agents are an important tool in the prevention of bacterial infections. Inorganic materials are attractive due to their high stability under a variety of conditions compared to organic antibacterial agents. Herein tungsten oxide nanodots (WO_3-X_), synthesized by a simple one-pot synthetic approach, was found to exhibit efficient antibacterial capabilities. The analyses with colony-forming units (CFU) showed excellent antibacterial activity of WO_3-X_ against both gram-negative *E. coli* (*Escherichia coli*) and gram-positive *S. aureus (Staphylococcus aureus*) strains. The scanning electron microscopy (SEM) and transmission electron microscopy (TEM) images revealed clear damage to the bacterial cell membranes, which was further confirmed by molecular dynamics simulations. Additionally, exposure to simulated sunlight was found to further increase germicidal activity of WO_3-X_ nanodots – a 30-minute exposure to sunlight (combining 50 μg/mL WO_3-X_ nanodots) showed a 70% decrease in *E. coli* viability compared to without exposure. Electron spin resonance spectroscopy (ESR) was used to elucidate the underlying mechanism of this photocatalytic activity through the generation of hydroxyl radical species. Cell counting kit-8 (CCK-8) and the live/dead assay were further employed to evaluate the cytotoxicity of WO_3-X_ nanodots on eukaryotic cells, which demonstrated their general biocompatibility. In all, our results suggest WO_3-X_ nanodots have considerable potential in antibacterial applications, while also being biocompatible at large.

## Introduction

Bacterial infections are a significant threat to a modern healthy population. Currently, close to 48 million cases of pathogenic diseases occur annually and approximately 88,000 people die from bacterial infection in the United States alone every year.^1-2^ Perhaps more troublesome, the abuse and overuse of antibiotics in recent decades have made them less effective against bacterial infections. This is specifically true for multidrug resistant bacteria aptly named “superbugs”. In the past few years, multidrug-resistance genes NDM-1 and MCR-1 were found in the plasmids of superbugs, which could rapidly transfer to other strains.^3-4^ Thus, new antibiotics that can effectively overcome the significant health threat of superbugs are urgently needed.

In the past few years, inorganic nanomaterials have drawn significant attention due to their distinctive properties of small size, large surface area, and high reactivity. Inorganic nanomaterials have been widely used in biomedical applications such as biocatalysis,^5-6^ cell imaging,^7-9^ drug delivery,^9-10^ tumor diagnosis and treatment.^11-12^ Moreover, inorganic nanomaterials have also been applied for antibacterial agents.^13-18^ Compared to organic materials, inorganic antibiotics are stable under harsh conditions such as high temperature and pressure.^19-21^ Due to high stability, inorganic materials have attracted higher levels of interest for bacterial control compared to organic antibiotics.^22-23^ A number of studies have demonstrated that some metal nanomaterials, such as Ag^13, 24-26^ and Cu^15, 27-28^, have strong germicidal activity. Unfortunately, sliver nanoparticles were found to be significantly toxic to mammalian cells due to the release of Ag^+^ ions and their interactions with thiol groups of proteins in mammalian cells.^29-30^ Additionally, copper nanoparticles were also found to cause serious injury to the kidney, liver, and spleen of exposed mice through oral gavage.^31^ Thus, significant biotoxicity has limited the widespread application of these metal nanoparticles.

Alternatively, metal oxide nanoparticles (such as TiO_2_ and ZnO) have been found to be more suitable for commercial bactericide agents because of their lower cost and less toxicity.^32-33^ In addition to antibacterial activity, many metal oxide materials exhibit photocatalytic properties.^34-36^ Under the stimulation of light, separation of charge could be induced within metal oxide materials resulting in the generation of electrons and holes. Due to the strong oxidative ability of holes, hydroxyl radicals (·OH) can be produced by the reaction of holes with water. Furthermore, electrons can reduce O_2_ generating super oxide anion (O_2_-_•_), another reactive oxygen species.^37-38^ Biologically, ·OH and O_2_-_•_ can cause peroxidation of the germ membrane lipids, strand breakage of DNA, inactivation of proteins, and eventually germ death.^39-41^

ZnO nanoparticles have been widely explored as a model metal oxide nanomaterial for its excellent bacterial-killing ability with or without irradiation.^42-46^ Unfortunately these materials have been found to be quite toxic. Chen and coworkers have elucidated that ZnO nanoparticles could activate endoplasmatic reticulum stress response and even induce apoptosis.^47^ Other *in vivo* studies have found ZnO nanoparticles can cause liver injury and induce hepatocyte apoptosis through oxidative stress and endoplasmatic reticulum stress.^48^ Therefore, more effort is needed in developing an effective antibacterial agent that not only has strong bactericidal ability, but is also biocompatible.

Previously, we have synthesized ultrasmall WO_3-X_ nanodots and explored their broad potential application for tumor theranostics and treatment.^49^ It was found that the antibacterial activity of WO_3-X_ nanodots was a result of membrane stress along with their photocatalytic properties, which provided a significant enhancement on their strong bactericidal activity. Both the scanning electron microscopy (SEM) and transmission electron microscopy (TEM) images clearly revealed damages to the bacterial cell membranes, which was further confirmed by our molecular dynamics (MD) simulations. Meanwhile, WO_3-X_ nanodots were found to be benign to eukaryotic cells. Our findings thus show that WO_3-X_ nanodots have significant potential as an effective, but biologically safe, antibacterial agent.

## Results and Discussion

### Synthesis and characterization of WO_3-X_

As shown in our previous study^49^, water-dispersible WO_3-X_ nanodots were synthesized by a simple one-pot reaction within 2h (**Figure 1**). TEM was used to study the morphology and size of WO_3-X_. As shown in **Figure 1a**, the WO_3-X_ nanodots were found to be uniform and ultra-small (1.1±0.3nm). Previous studies have demonstrated that smaller particles were more effective in killing germs as compared to larger particles. It has been proposed that smaller nanoparticles may penetrate the cell walls of bacteria and interact with membranes more easily.^42, 50^

**Figure 1.**
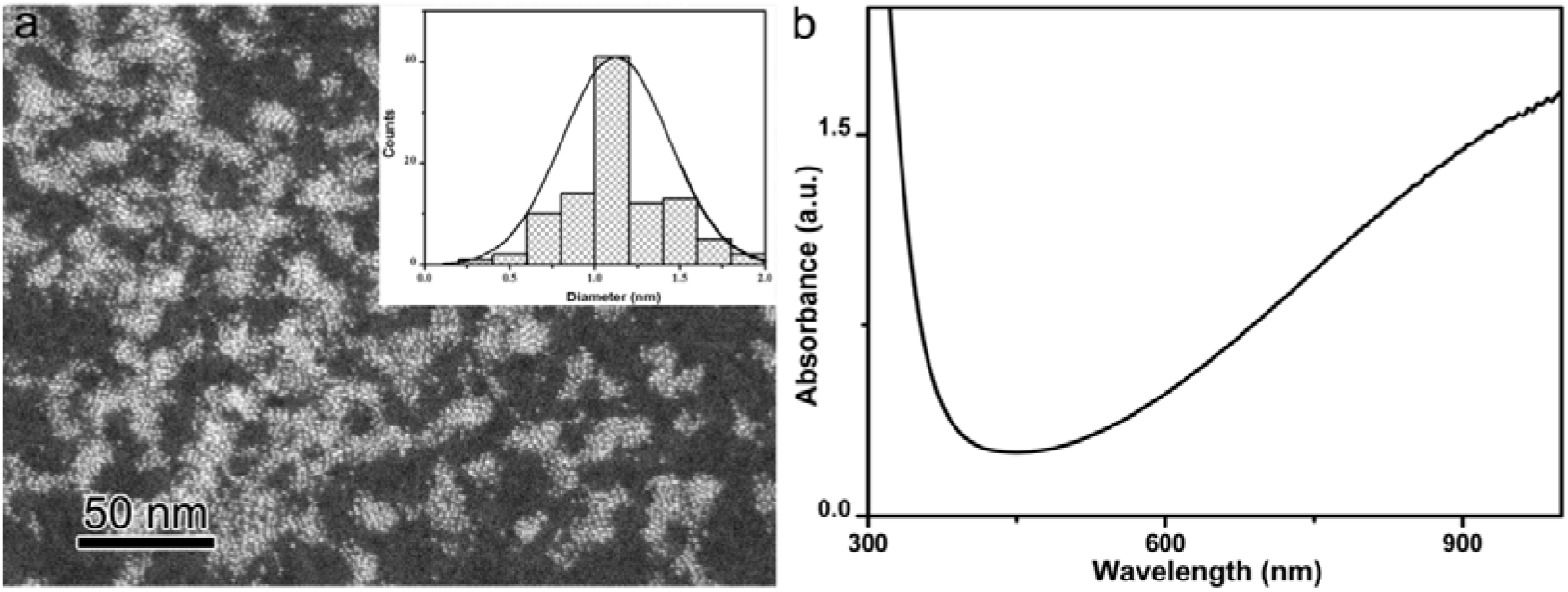
Characterization of WO_3-X_ nanodots. (a) TEM imaging and size distribution (inset) of WO_3-X·_ (b) UV-absorption spectrum of WO_3-X_ nanodots.

UV-vis spectrum of the WO_3-X_ nanodots exhibited a strong absorption band across the near-infrared range (NIR), making them an ideal probe for photothermal therapy and photoacoustic imaging of tumors. Our previous study focused on the application of WO_3-X_ nanodots for cancer theranostics.^49^ Additionally, we have previously used X-ray diffraction (XPS) and X-ray photoelectron spectroscopy (XRD) to analyze the structure of WO_3-X_. We found the nanodots were oxygen deficient and their crystalline structure were similar to the nonstoichimetric monoclinic WO_2.72_.^49^

### Antibacterial activity

To evaluate the antibacterial activity of WO_3-X_ nanodots, a standard colony counting assay was employed. As demonstrated in **Figure 2**, after 2h of treatment with less than 50 μg/mL of WO_3-X_, slight germicidal activity to *E. coli* was detected. The viability of *E. coli* cells were 95.7% and 84.6% corresponding to 25μg/mL and 50 μg/mL treatment concentration respectively. Whereas for the treatment concentration up to 100μg/mL, cell viability decreased to 19.7%, which illustrated that WO_3-X_ could kill bacteria in a concentration-dependent manner. Moreover, with the increase of exposure time, the antibacterial activity of WO_3-X_ also increased significantly. Viability of *E. coli* decreased to 44.0%, 37.0% and 0%, when treated with 25, 50 and 100μg/mL WO_3-X_ for 6h. A similar trend was also found in *S. aureus*. However, compared to *E. coli*, *S. aureus* was more sensitive to WO_3-X_. It was found that 25, 50 and 100μg/mL WO_3-X_ could kill 11.1%, 48.8% and 88.9% *S. aureus* when treated for 2 h and loss of viability increased to 81.3%, 90.0%, 100%, respectively, when treated for 6h. Previous studies attributed different sensitivity to antibacterial agents between Gram-positive and Gram-negative bacterial strains to their differences in the structure of cell walls and membranes.^51-52^ Gram-negative bacteria possess an outer membrane which is not present in gram-positive bacteria. An additional membrane may induce the resistance of gram-negative bacteria to WO_3-X_ by providing another physical layer.

**Figure 2.**
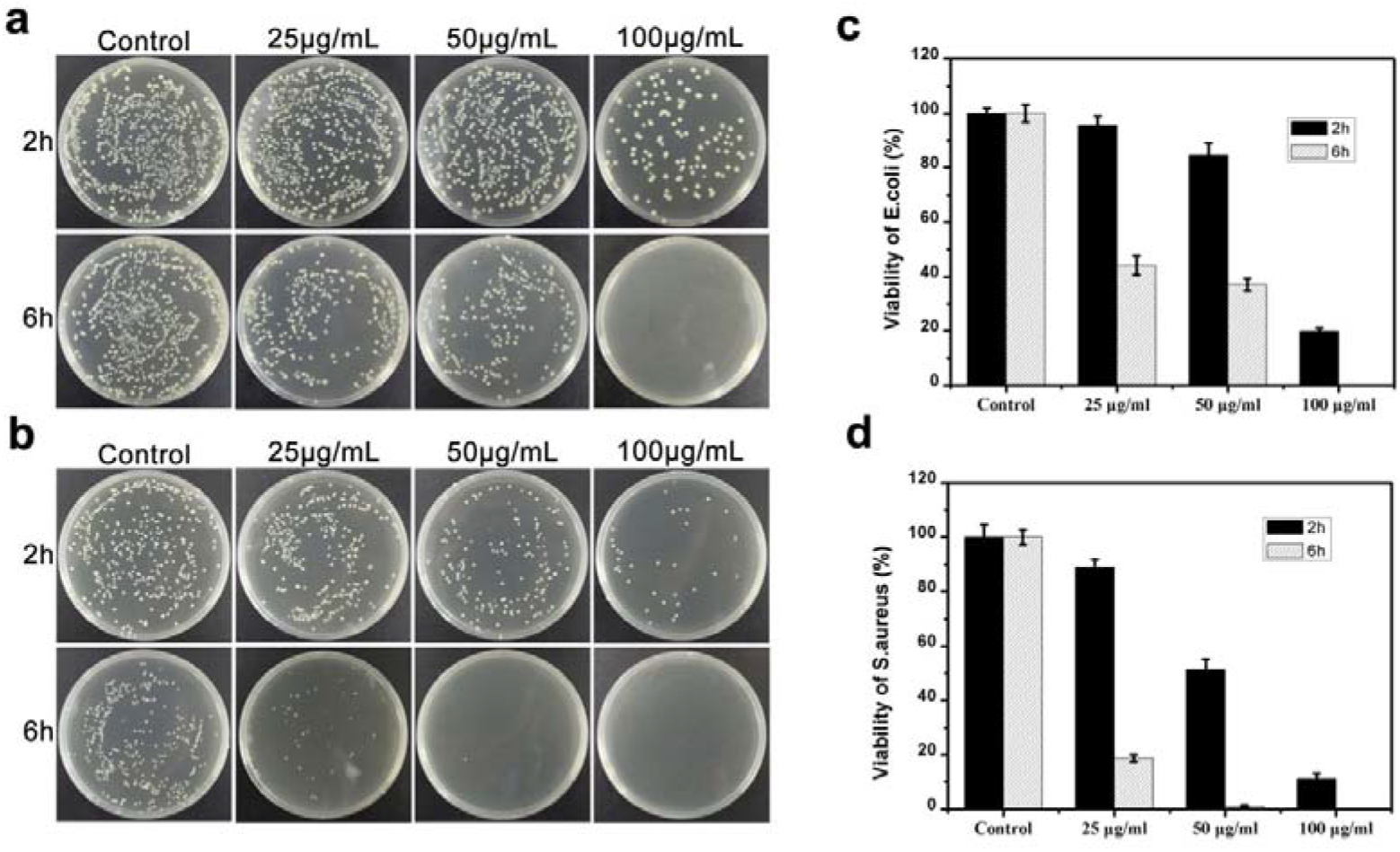
Antibacterial activity of WO_3-X_ nanodots. (a,c) viability of *E. coli* and *S. aureus* (b,d) after treated with different concentrations of WO_3-X_ nanodots for 2 and 6h.

### Bacterial morphology changes

To further investigate the antibacterial effect of WO_3-X_, SEM was used to investigate the morphology changes caused by WO_3-X_. As depicted in **Figure 3**, both *E. coli* and *S. aureus* cells were well defined and fully intact with the WO_3-X_ absent. Conversely, after incubation with 100μg/mL WO_3-X_ for 2h, a significant amount of WO_3-X_ adhered onto the surface of the bacterial cells. Thus, the bacterial cells were found to deform significantly. SEM revealed the surface of *E. coli* cells became coarse and potholed. Correspondingly, most *S. aureus* cells were found to collapse suggesting loss of their cytoplasm. Previous studies found that numerous nanomaterials could kill bacteria due to direct interaction with the membrane leading to physical disruption.^43, 53-55^

**Figure 3.**
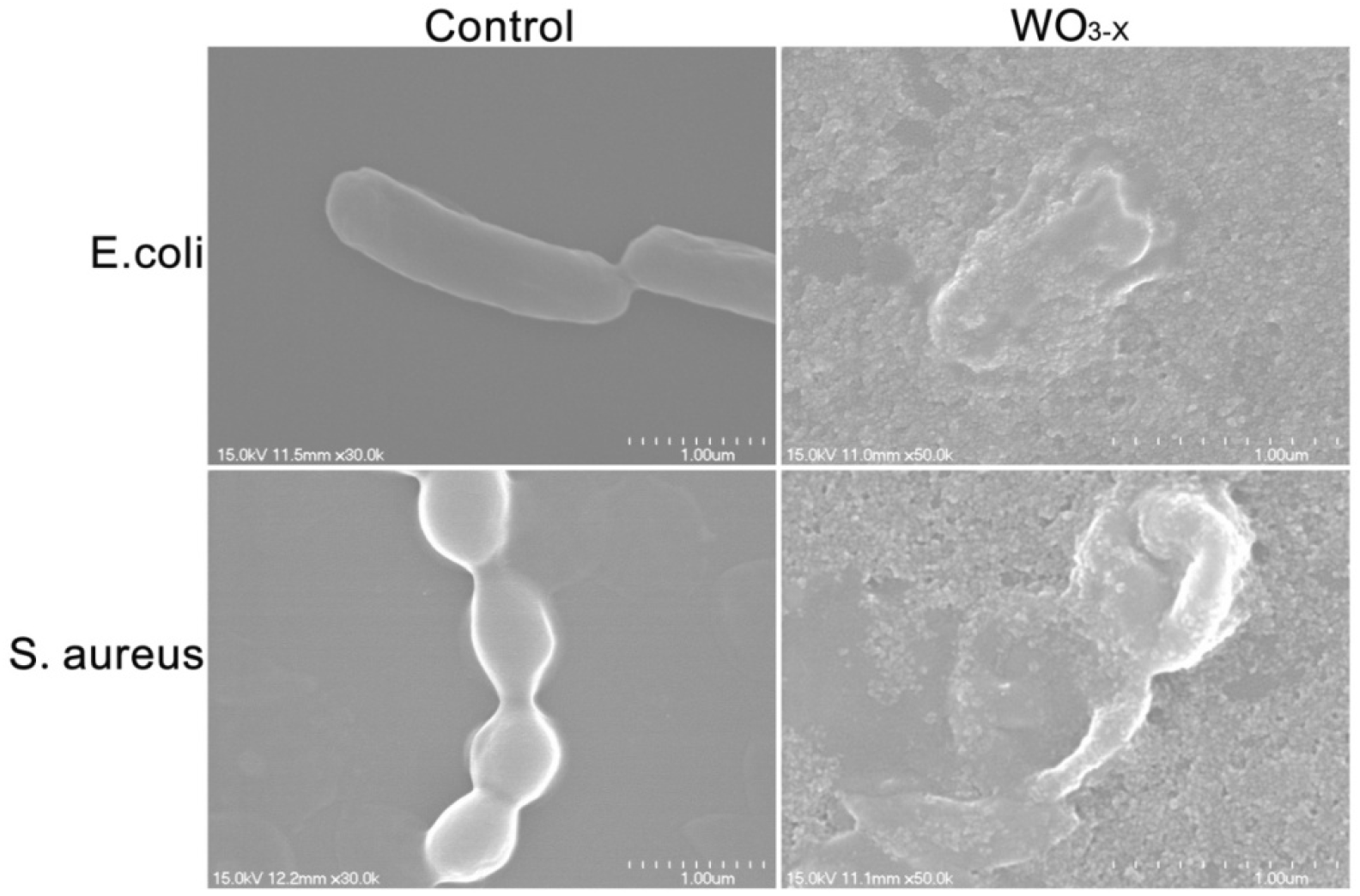
Morphological damage of *E. coli* and *S. aureus* was detected by SEM imaging after treated with 100μg/mL WO_3-X_ for 2h.

Moreover, TEM imaging was also employed to observe the mechanism by which WO_3-X_ impaired bacteria. Morphologies of normal *E. coli* cells (control) were shown as regular and integrated, with uniform cytoplasm (Figure 4; left). However, significant damage of the *E. coli* membranes were observed after the incubation of 100μg/mL WO_3-X_ for 2h (Figure 4; right). The location of WO_3-X_ nanodots on the *E. coli* membrane surface were marked with red arrows (Figure 4; right). These additional TEM images clearly demonstrate the severe injury of these bacterial cells, with both damage in membrane and effusion of cytoplasm, consistent with the findings from the SEM imaging.

**Figure 4.**
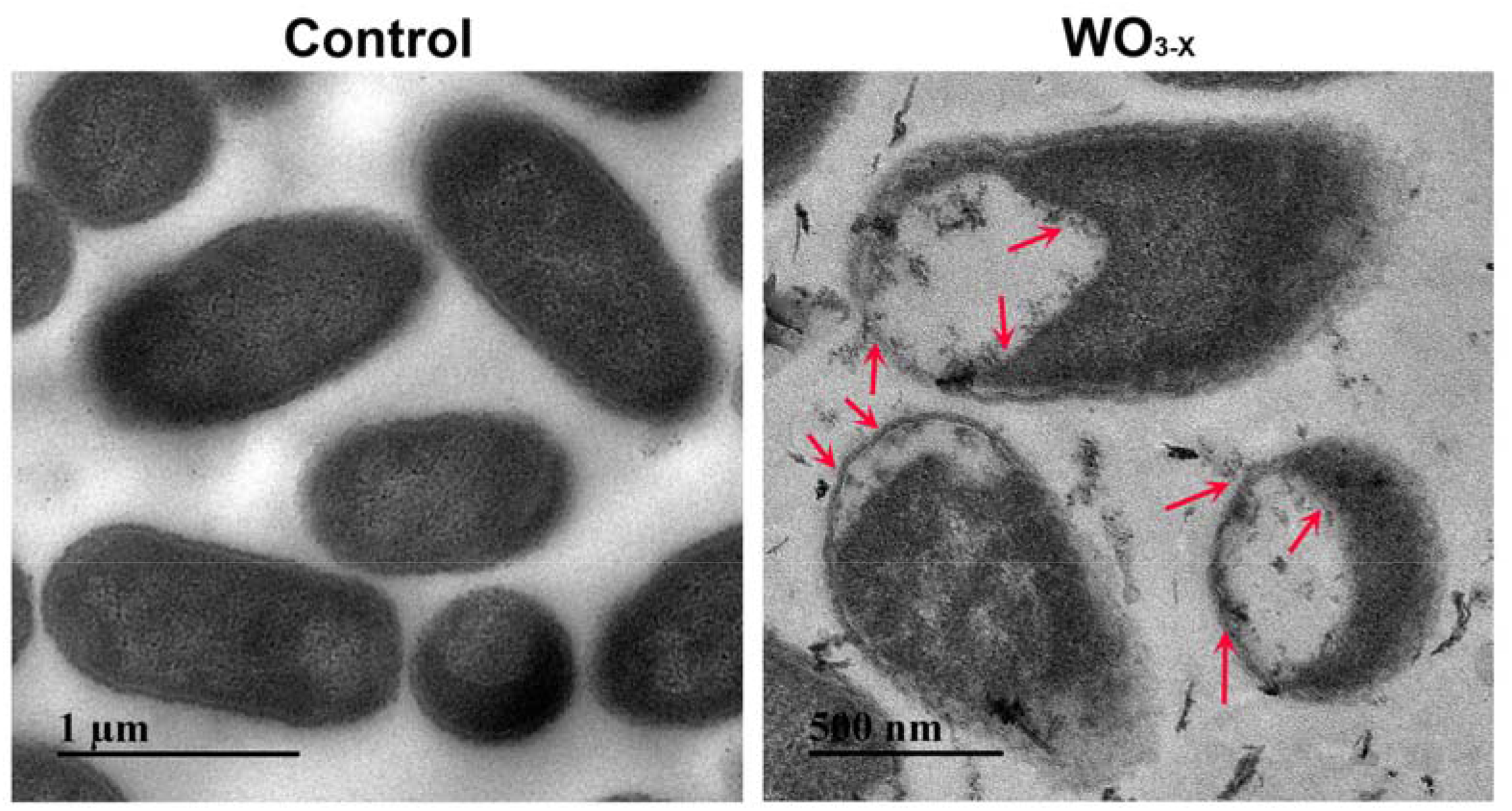
Membrane impairment of *E. coli* after the treatment of 100μg/mL WO_3-X_ for 2h with TEM imaging (left: control; right: with WO_3-X_). The location of WO_3-X_ nanorods on the membrane surfaces were marked by red arrows.

The above antibacterial assays, along with SEM and TEM images suggest that WO_3-X_ might interact with the membrane of bacterium directly and disturb the function of membrane.

### Interaction between WO_3-X_ and bacterial membrane

Molecular dynamics (MD) simulations were used to probe the interaction mechanism between WO_3-X_ and the bacterial membrane (see Figure 5). Figure 5a shows the initial conformation of the membrane and nanodots system, and Figure 5b shows a representative conformation during the unbiased MD simulations. The nanodots reversibly adsorb onto the membrane within 50 ns, leading to accumulation of WO_3-X_ near the membrane (Figure 5d). They can either bind directly to the membrane or indirectly interact through a layer of bridging water. No permeation into the membrane was observed during 150 ns of simulation time. Energy analysis (Figure 5c) shows that the binding between membrane and WO_3-x_ is mainly driven by Coulomb interactions. Additional “docking” simulations of the membrane and nanosheet system shows that the nanosheet can attach to the membrane but does not extract lipid molecules during 150 ns of simulations (Figure S3). This is in contrast to graphene nanosheets which can robustly extract lipid molecules from the membrane^56^. Taken together, the MD simulations indicate that WO_3-x_ interacts strongly with the lipid head groups and adsorb onto the surface of the membrane. The adsorption of WO_3-X_ could directly interfere with the functions of membrane and membrane proteins or enhance the activity of reactive oxygen species.

**Figure 5.**
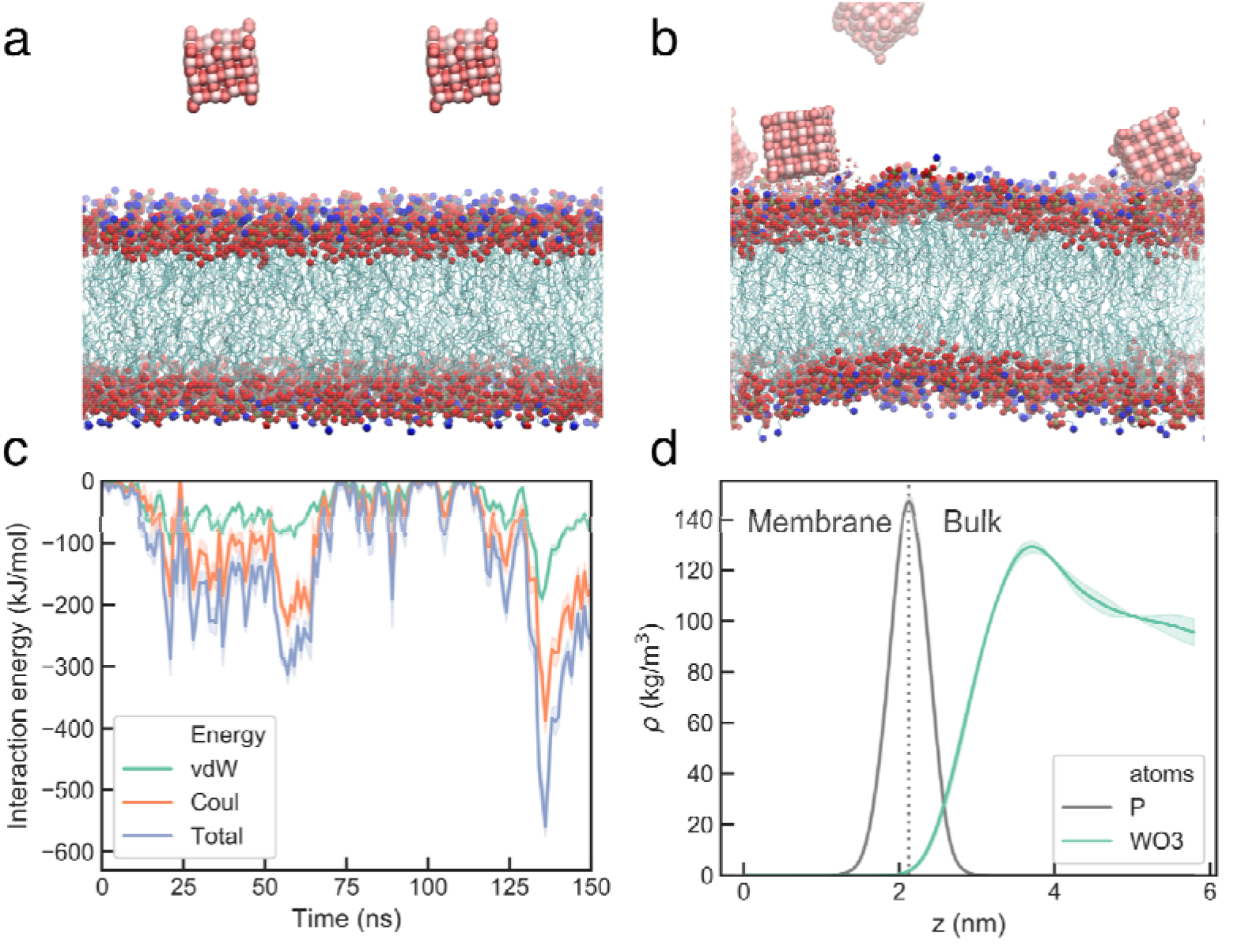
Molecular dynamics simulations of bacterial membrane with WO_3-X_ nanodots. (a) Starting conformation and (b) representative conformation at 60 ns. W, O, N and P atoms are shown in spheres, and the fatty acid tails are shown in sticks. (c) evolution of interaction energy between membrane and WO_3-X_ during the 150ns simulation. The energy is averaged over 1-ns windows. (d) Density distribution of WO_3-X_ along the normal direction to the membrane. The width of the band shows statistical error estimated from two independent simulations. The WO_3-X_ nanodots reversibly adsorb on the membrane surface due to interaction with phosphate and amonium groups on the lipids.

### Photoinduced antibacterial activity of WO_3-X_ nanodots

To further explore the antibacterial applications of WO_3-X_, photoinduced antibacterial activity was investigated (**Figure 6**). Our results showed that simulated sunlight exhibited limited toxicity to *E. coli*. As a control we found that even after being irradiated for 30 min (without WO_3-X_) survival rate was high (~88%; **Figure 6a-b**). However, after the bacteria were combined with WO_3-X_ in the presence of simulated sunlight, bacterial survival rate decreased significantly (**Figure 6a-b**). When *E. coli* was exposed to 50μg/mL WO_3-X_ nanodots and irradiated for 10 min, the viability decreased to ~21.2%. When irradiation time increased to 30 min, only 7.1% of *E. coli* cells survived, which was much less than the nonirradiated control (80.8%). To investigate the mechanism of photoinduced antibacterial activity of WO_3-X_, photoinduced ROS generation ability was measured. Previously, it was shown that ESR spectroscopy was a reliable method for the analysis of ROS generation.^57-58^ 5-tert-butoxycarbonyl 5-methyl-1-pyrroline N-oxide (BMPO), a spin trap for hydroxyl radical and superoxide was also required in this assay. As shown in **Figure 6c**, no ESR signal was detected in the control groups (consisted of unirradiated WO_3-X_ sample, irradiated WO_3-X_ sample without BMPO and WO_3-X_ absent sample). However, characteristic four-line spectrum with the relative intensities of 1:2:2:1 was detected after WO_3-X_ was exposed to simulated sunlight, which indicated that hydroxyl radical was generated by WO_3-X_ under irradiation. Moreover, signal intensity of the spectrum increased with the increase of irradiation time. These results suggest that the photo-induced antibacterial activity of WO_3-X_ nanodots can be attributed to photoexcited ROS generation. In addition, many studies have reported the mechanisms of photocatalytic activity of semiconductors. As a semiconductor, WO_3-X_ has a wide band gap and exhibits photocatalytic properties. Numerous studies have demonstrated that after semiconductor materials are irradiated with light such that the absorbed energy is equivalent to band gap, electrons in valence band will be excited to the conduction band resulting in the generation of hole and electron charge carriers.^35, 38, 44^ As a result of the strong oxidative ability of holes, hydroxyl radicals may be produced by reacting with water which is another ROS species that may contribute to antibacterial activity.

**Figure 6.**
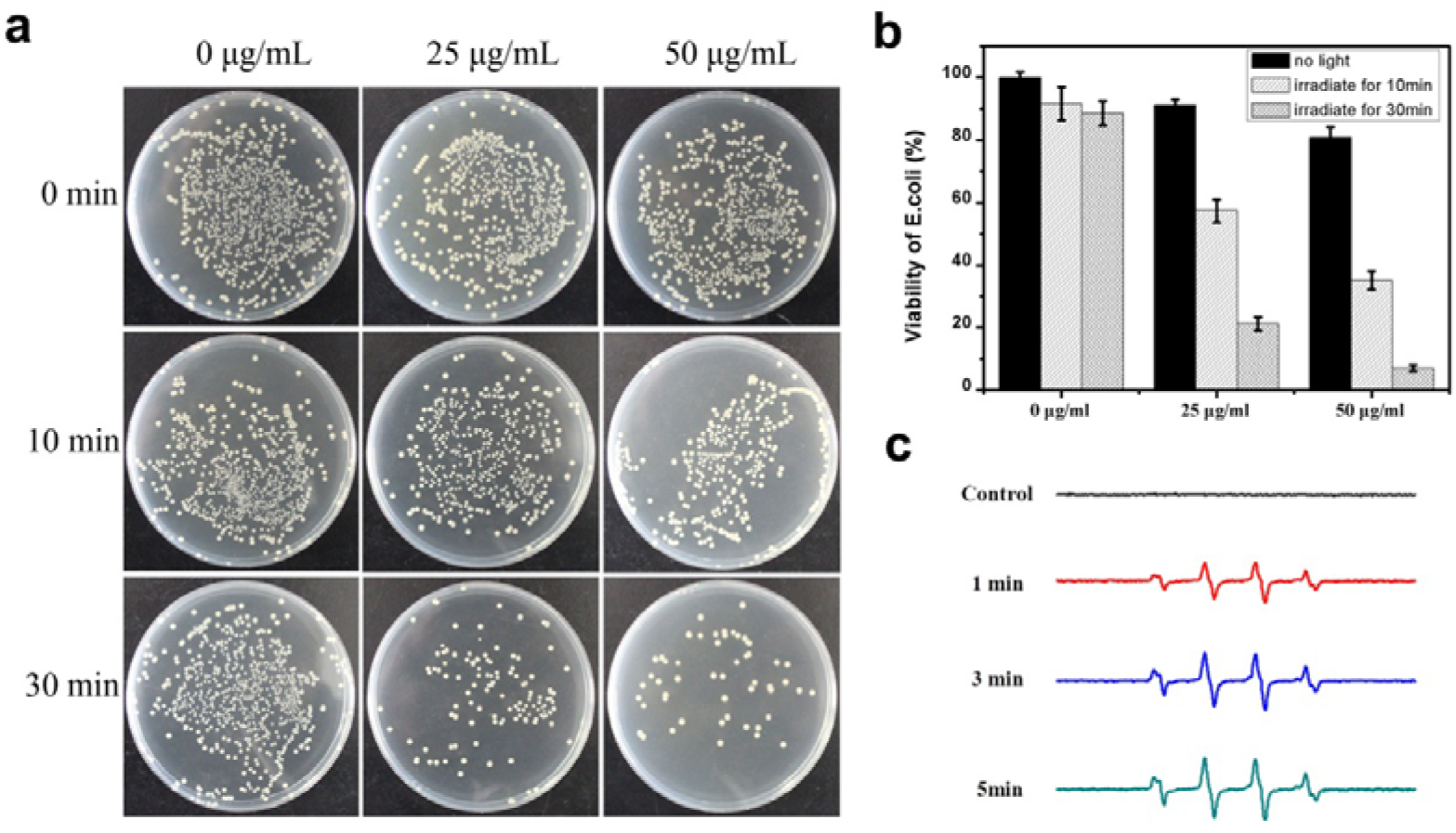
Photoinduced Antibacterial activity and photocatalytic activity of WO_3-X_ against *E. coli*. (a,b) Antibacterial activity of WO_3-X_ nanodots under simulated sunlight. (c). Photoinduced ROS generation ability of WO_3-X_ nanodots was detected by ESR spectra.

### Cytotoxicity to mammalian cells (Biocompatibility)

Because of its obvious antibacterial effect, cytotoxicity of WO_3-X_ to eukaryotic cells was also investigated. Human bronchial epithelial cell line (Beas-2B) and human umbilical vein endothelial cell (HUVEC) were chosen to evaluate cytotoxicity caused by WO_3-X_. Both CCK-8 and live/dead assays were implemented after different concentrations of WO_3-X_ were co-incubated with cells for 24h. As indicated in **Figure 7**, Beas-2b (7a) and HUVEC (7b) cells were not sensitive to WO_3-X_. Cell viability was still more than 80% even when cells were exposed to 400μg/mL of WO_3-X_. The results of live/dead assay (7c) also demonstrated that WO_3-X_ was non-toxic to Beas-2b and HUVEC cells.

**Figure 7.**
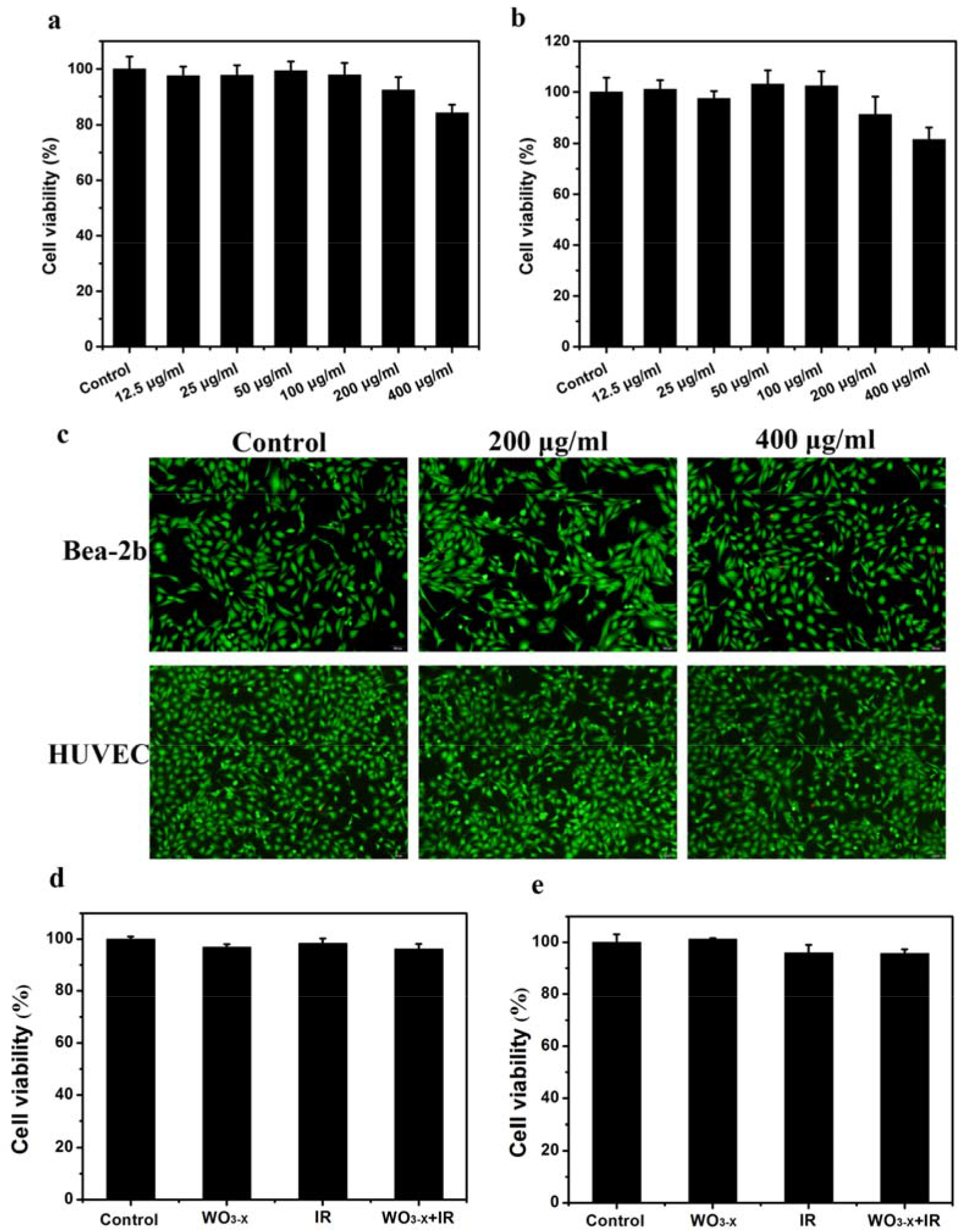
Cytotoxicity of WO_3-X_ nanodots to eukaryotic Beas-2b and HUVEC cells. Cell viability of Beas-2b (a) and HUVEC (b) after treated with different concentrations of WO_3-X_ nanodots for 24h was analyzed by CCK-8 and live/dead assay (c). Cytotoxicity of WO_3-X_ nanodots to Beas-2b (d) and HUVEC (e) cells combining simulated sunlight.

In addition, the cell viability of Beas-2b and HUVEC cells incubated with WO_3-X_ in combination of simulated sunlight was also analyzed by CCK-8 assay. After being treated with WO_3-X_ nanodots (200μg/mL) for 1 h and then closed with simulated sunlight exposure for 30 min, these eukaryotic cells were incubated for 24h to examine their cell viability. Different from bacteria, both Beas-2B (Figure 7d) and HUVEC (Figure 7e) were not sensitive to the photocatalytic effect of WO_3-X_. Simulated sunlight could not intensify the cytotoxicity of WO_3-X_, which might result from the auto antioxidant system in mammalian cells as compared to bacteria. The auto antioxidant system includes many antioxidant enzymes, such as superoxide dismutase (SOD), peroxidase (POD), catalase (CAT) and so on^59-60^. Those enzymes could relieve the oxidative stress caused by WO_3-X_ nanodots when exposed to simulated sunlight.

The difference of sensitivity to WO_3-X_ between bacteria and eukaryotic cells may be attributed to the difference in their culture environment. Because of their large surface area and high surface free energy, nanomaterials may adsorb biomolecules (such as fetal bovine serum proteins in complete medium) onto their surfaces and form a surface coating structure (the so-called protein ‘corona’).^61^ Corona surface structures may lower the surface free energy of nanomaterials and changes their dispersibility.^61^ In addition, corona has also been proved to change the surface properties of nanomaterials and alter their biological responses.^61-62^ For example, in our previous studies, we have indicated that fetal bovine serum proteins can absorb easily onto graphene oxide (GO) nanosheets to form corona, which weakens the GO-phospholipids interaction and results in a less destructive action to cell membranes due to the unfavorable steric effects of corona on GO surfaces.^62-63^ Furthermore, the auto antioxidant system in mammalian cells may help reduce the oxidative stresses.

### Conclusion

In summary WO_3-X_ nanodots were synthesized by a one-pot approach within 2h and subsequently tested as an antibacterial agent. The antibacterial activity of WO_3-X_ nanodots were investigated against both Gram-negative *E. coli* and Gram-positive *S. aureus*. It was found that as-produced WO_3-X_ nanodots could kill both *E. coli* and *S. aureus* in a concentration- and time-dependent manner. Our findings further suggested that the antibacterial mechanism of WO_3-X_ may be attributed to bacterial membrane disruption. Additionally, we also detected that simulated sunlight could significantly increase the antibacterial activity of WO_3-X_ nanodots by inducing the generation of reactive oxygen species (ROS). Importantly, WO_3-X_ nanodots were found to be nontoxic to eukaryotic cells even combining the simulated sunlight, which suggesting far reaching potential as a selective antibacterial agent.

## Experimental section

### Synthesis and characterization of WO_3-X_

WO_3-X_ was synthesized by a simple and rapid reaction, which was described in detail from our previous study.^49^ Briefly, 80 mL of pentaerythritol tetrakis (3-mercaptopropionate)-terminated poly(methacrylic acid) (PTMP-PMAA) solution was heated to boil under a nitrogen environment. 20 mL WCl6 was then added and reacted for 2h. The mixture was then cooled and centrifuged with an ultrafiltration tube to remove residual PTMP-PMAA. WO_3-X_ was characterized with transmission electron microscopy (TEM) and an ultraviolet-visible (UV-vis) spectrophotometer. The WO_3-X_ solution was casted onto a copper grid and then dried before characterization. TEM images were collected by using a FEI Tecnai G20 TEM with an acceleration voltage of 200kV. UV-vis absorption spectrum was recorded by a PerkinElmer Lambda 750 UV-vis-NIR spectrophotometer.

### Bacterial culture and antibacterial activity assay

*E. coli* (ATCC25922,) and *S. aureus* (ATCC25923,) were selected as representative gram-negative and gram-positive bacterial models, respectively, for the antibacterial assays. Both types of bacterium were grown in LB (Luria-Bertani) medium at 37 °C overnight and harvested at the mid-exponential growth phase. Next, the bacteria were washed with isotonic saline solution 3 times to rinse away any residual medium. Subsequently, bacterial cells were diluted to the concentration of 10^6^ to 10^7^ colony forming units per milliliter (CFU/mL) with isotonic saline solution. To examine the antibacterial activity, 0, 25, 50, 100 μg/mL WO_3-X_ nanodots were used to treat both *E. coli* and *S. aureus* for 2h and 6h under the condition of 37°C and 250 rpm shaking. Finally, 100 μL bacterium solution was spread on LB plates and after an overnight incubation, colonies were counted to evaluate antibacterial activity of WO_3-X_ to both *E. coli* and *S. aureus*.

### Bacterial morphology detection

To visualize the morphological changes of *E. coli* and *S. aureus* bacterial cells after WO_3-X_ treatment, we incubated both types of bacteria with WO_3-X_ at the final concentration of 100 μg/mL at 37 °C and 250 rpm shaking for 2h. The bacteria were then centrifuged and washed with sterile deionized water 3 times. Next, bacterial cells were prefixed with 2.5% glutaraldehyde for 2h and fixed with 1% osmium tetroxide for 1h sequentially. Subsequently, dehydration was executed by using a graded series of ethanol (30%, 50%, 70%, 80%, 90%, 95% and 100%). Finally, bacterial cells were dried in a vacuum oven, sputter-coated with gold and imaged under the field-emission SEM. TEM samples were prepared following the previous study^62^. Briefly, after being incubated with WO_3-X_ nanodots, *E.coli* cells were fixed with 2.5% glutaraldehyde overnight. Then samples were fixed again with OsO4, dehydrated with gradient ethanol solution, embedded with resin, and then cut into sections of 70 nm in thickness. Finally, sections were loaded onto copper grids and scanned by HT7700 transmission electron microscopy (Hitachi).

### Sunlight induced ROS generation and antibacterial activity analysis

ESR was used to observe the photocatalytic activity of WO_3-X_. The spin-trap 5-tert-butoxycarbonyl 5-methyl-1-pyrroline N-oxide (BMPO) was employed to verify the generation of ·OH radical species. 50μL of WO_3-X_ (100μg/mL) was loaded into quartz capillary tubes and mixed with BMPO (the final concentration was 25mM). The capillary tubes were sealed and inserted into the ESR cavity. Spectra signals were recorded when samples were exposed to simulated sunlight (provide by filtered 450 W Xenon light) for selected times of 0, 1, 3 and 5 minutes.

To investigate the antibacterial activity of WO_3-X_ after exposure to sunlight, the mixtures including *E. coli* and different concentrations of WO_3-X_ were irradiated under a simulated sunlight for 0, 10 and 30 min. After incubation for a total treatment time of 2h at 37°C, 100μL mixtures were spread onto LB plates. Next, plates were incubated at 37°C overnight. Resultant colonies on the plates were counted to calculate survival rate of bacterium.

### Cytotoxicity assay

Beas-2B cells and HUVEC cell lines were used to investigate cytotoxicity of WO_3-X_. Beas-2B and HUVEC cells were cultured with DMEM medium supplied with 10% fetal bovine serum (FBS) and 1% antibiotic (streptomycin 100μg/mL and penicillin 100 U/mL) in an incubator at 37°C and in a 5% CO2 environment. Cells were seeded in 96-well plates at the concentration of 5000 cells/well. Following a 24h incubation, cells were incubated with fresh medium containing 0, 25, 50, 100, 200, 400μg/mL WO_3-X_. After treatment for 24h, cell counting kit-8 (CCK-8) was added in each well. The cells were then incubated for an additional 2h at 37 °C and 5% CO_2_. The cytotoxicity of WO_3-X_ was analyzed by measuring spectrophotometrical absorbance at 450 nm wavelength by a microplate reader.

### Molecular dynamics simulations

Molecular dynamics simulations were performed to understand the interaction between WO_3-X_ and the bacteria cell membrane. The CHARMM36m force field was used to model lipids and water. For WO_3-X_, the partial charge on oxygen was determined from quantum mechanics (QM) density functional theory (DFT) calculations and Mulliken and Lowdin analyses (Figure S1, Table S1), and then the vdW parameters were tuned to reproduce the experimental water contact angle (ca. 0 degree) and QM equilibrium interaction distance between water and WO_3-X_ (Figure S2, Table S2). The bond lengths and angles were taken from the crystal structure. The WO_3-X_ nanodot model was constructed by cutting four layers of the W18O49 crystal structure and has the stoichiometry of W_48_O_112_ (i.e. x=0.67 in the reduced formula WO_3-X_) The bacteria membrane was represented by a lipid bilayer consisting of 342 POPE molecules.

Following similar protocols in our previous studies^64-76^, four WO_3-X_ nanodots and the membrane were placed in a simulation box with dimension 9.9×9.9×12.0 nm^3^. The nanodots were at least 2 nm away from the membrane in the initial structure. Multiple 150 ns long MD simulations were performed for the complex system. Additional “docking” simulations were also performed in which a WO_3-X_ nanosheet was placed perpendicularly to the membrane. The nanosheet with a size of 4.1×1.2×7.0 nm^3^ was constructed by cutting four layers of the W18O49 crystal structure and removing surface oxygen atoms. The size of the simulation box was 9.8×9.8×16.8 nm^3^. The initial separation distance between the nanosheet and the membrane was 1 nm. Then the positions of the nanosheet were fixed while the membrane was fully flexible.

## Supporting information

## Acknowledgements

The authors gratefully acknowledge the help from Zonglin Gu and Tien Huynh. This work was partially supported by the National Natural Science Foundation of China under Grant Nos. 11374221, 11574224, and 21320102003. RZ acknowledges the support from IBM Blue Gene Science Program (W125859, W1464125, W1464164). A project partly funded by the Priority Academic Program Development of Jiangsu Higher Education Institutions (PAPD), Jiangsu Provincial Key Laboratory of Radiation Medicine and Protection, Suzhou Scientific and Technology Program (sys2018087), and Suzhou Project of Health Development through Science & Education (KJXW2017039).

## Author contributions

R.Z., G.D. and J.X. conceived and designed the experiments. G.D., L.C., L.Z., and J.X., performed the antibacterial experiments. Z.J. and R.Z. performed the molecular dynamics simulations. L.W. and Z.L. synthesized the nanomaterials. G.D., L.C., R.Z., Z.J., P.D.L., L.Z., J.X. and Z.Y. analyzed the data. G.D., Z.J., P.D.L.,Y.Z. and R.Z., co-wrote the paper. All authors discussed the results and commented on the manuscript.

## Additional information

The authors declare no competing financial interests. Correspondence and requests for materials should be addressed to R.H.Z.

## References

1. Morris, J. G., How Safe Is Our Food? Emerg. Infect. Dis 2011, 17, 126–128.

2. Hsueh, P. R.; Huang, H. C.; Young, T. G.; Su, C. Y.; Liu, C. S.; Yen, M. Y., Bacteria Killing Nanotechnology Bio-Kil Effectively Reduces Bacterial Burden in Intensive Care Units. European Journal of Clinical Microbiology & Infectious Diseases 2014, 33, 591–597.

3. Yong, D.; Toleman, M. A.; Giske, C. G.; Cho, H. S.; Sundman, K.; Lee, K.; Walsh, T. R., Characterization of a New Metallo-Beta-Lactamase Gene, Bla(Ndm-1), and a Novel Erythromycin Esterase Gene Carried on a Unique Genetic Structure in Klebsiella Pneumoniae Sequence Type 14 from India. Antimicrob. Agents Chemother. 2009, 53, 5046–5054.

4. Zhang, R.; Huang, Y. L.; Chan, E. W. C.; Zhou, H. W.; Chen, S., Dissemination of the Mcr-1 Colistin Resistance Gene. Lancet Infect. Dis. 2016, 16, 291–292.

5. Yang, H. H.; Zhang, S. Q.; Chen, X. L.; Zhuang, Z. X.; Xu, J. G.; Wang, X. R., Magnetite-Containing Spherical Silica Nanoparticles for Biocatalysis and Bioseparations. Anal. Chem. 2004, 76, 1316–1321.

6. Xie, W. L.; Ma, N., Immobilized Lipase on Fe3o4 Nanoparticles as Biocatalyst for Biodiesel Production. Energy Fuels 2009, 23, 1347–1353.

7. Huang, X. H.; El-Sayed, I. H.; Qian, W.; El-Sayed, M. A., Cancer Cell Imaging and Photothermal Therapy in the near-infrared Region by Using Gold Nanorods. J. Am. Chem. Soc. 2006, 128, 2115–2120.

8. Santiago, A. M.; Ribeiro, T.; Rodrigues, A. S.; Ribeiro, B.; Frade, R. F. M.; Baleizao, C.; Farinha, J. P. S., Multifunctional Hybrid Silica Nanoparticles with a Fluorescent Core and Active Targeting Shell for Fluorescence Imaging Biodiagnostic Applications. Eur. J. Inorg. Chem. 2015, 4579–4587.

9. Liong, M.; Lu, J.; Kovochich, M.; Xia, T.; Ruehm, S. G.; Nel, A. E.; Tamanoi, F.; Zink, J. I., Multifunctional Inorganic Nanoparticles for Imaging, Targeting, and Drug Delivery. ACS Nano 2008, 2, 889–896.

10. Chen, Y.; Chen, H. R.; Zhang, S. J.; Chen, F.; Zhang, L. X.; Zhang, J. M.; Zhu, M.; Wu, H. X.; Guo, L. M.; Feng, J. W., et al., Multifunctional Mesoporous Nanoellipsoids for Biological Bimodal Imaging and Magnetically Targeted Delivery of Anticancer Drugs. Adv. Funct. Mater. 2011, 21, 270–278.

11. Yezhelyev, M. V.; Gao, X.; Xing, Y.; Al-Hajj, A.; Nie, S. M.; O’Regan, R. M., Emerging Use of Nanoparticles in Diagnosis and Treatment of Breast Cancer. Lancet Oncol. 2006, 7, 657–667.

12. Cuenca, A. G.; Jiang, H. B.; Hochwald, S. N.; Delano, M.; Cance, W. G.; Grobmyer, S. R., Emerging Implications of Nanotechnology on Cancer Diagnostics and Therapeutics. Cancer 2006, 107, 459–466.

13. Mohan, Y. M.; Lee, K.; Premkumar, T.; Geckeler, K. E., Hydrogel Networks as Nanoreactors: A Novel Approach to Silver Nanoparticles for Antibacterial Applications. Polymer 2007, 48, 158–164.

14. Banoee, M.; Seif, S.; Nazari, Z. E.; Jafari-Fesharaki, P.; Shahverdi, H. R.; Moballegh, A.; Moghaddam, K. M.; Shahverdi, A. R., Zno Nanoparticles Enhanced Antibacterial Activity of Ciprofloxacin against Staphylococcus Aureus and Escherichia Coli. Journal of Biomedical Materials Research Part B-Applied Biomaterials 2010, 93B, 557–561.

15. Longano, D.; Ditaranto, N.; Cioffi, N.; Di Niso, F.; Sibillano, T.; Ancona, A.; Conte, A.; Del Nobile, M. A.; Sabbatini, L.; Torsi, L., Analytical Characterization of Laser-Generated Copper Nanoparticles for Antibacterial Composite Food Packaging. Analytical and Bioanalytical Chemistry 2012, 403, 1179–1186.

16. Subbiandoss, G.; Sharifi, S.; Grijpma, D. W.; Laurent, S.; van der Mei, H. C.; Mahmoudi, M.; Busscher, H. J., Magnetic Targeting of Surface-Modified Superparamagnetic Iron Oxide Nanoparticles Yields Antibacterial Efficacy against Biofilms of Gentamicin-Resistant Staphylococci. Acta Biomaterialia 2012, 8, 2047–2055.

17. Zhao, Y. Y.; Ye, C. J.; Liu, W. W.; Chen, R.; Jiang, X. Y., Tuning the Composition of Aupt Bimetallic Nanoparticles for Antibacterial Application. Angewandte Chemie-International Edition 2014, 53, 8127–8131.

18. Paul, T.; Miller, P. L.; Strathmann, T. J., Visible-Light-Mediated Tio2 Photocatalysis of Fluoroquinolone Antibacterial Agents. Environ. Sci. Technol. 2007, 41, 4720–4727.

19. Makhluf, S.; Dror, R.; Nitzan, Y.; Abramovich, Y.; Jelinek, R.; Gedanken, A., Microwave-Assisted Synthesis of Nanocrystalline Mgo and Its Use as a Bacteriocide. Adv. Funct. Mater. 2005, 15, 1708–1715.

20. Sawai, J.; Igarashi, H.; Hashimoto, A.; Kokugan, T.; Shimizu, M., Effect of Particle Size and Heating Temperature of Ceramic Powders on Antibacterial Activity of Their Slurries. Journal of Chemical Engineering of Japan 1996, 29, 251–256.

21. Hewitt, C. J.; Bellara, S. R.; Andreani, A.; Nebe-von-Caron, G.; McFarlane, C. M., An Evaluation of the Anti-Bacterial Action of Ceramic Powder Slurries Using Multi-Parameter Flow Cytometry. Biotechnology Letters 2001, 23, 667–675.

22. Tang, Z. X.; Lv, B. F., Mgo Nanoparticles as Antibacterial Agent: Preparation and Activity. Brazilian Journal of Chemical Engineering 2014, 31, 591–601.

23. Sevinc, B. A.; Hanley, L., Antibacterial Activity of Dental Composites Containing Zinc Oxide Nanoparticles. Journal of Biomedical Materials Research Part B-Applied Biomaterials 2010, 94B, 22–31.

24. Kim, J. S.; Kuk, E.; Yu, K. N.; Kim, J. H.; Park, S. J.; Lee, H. J.; Kim, S. H.; Park, Y. K.; Park, Y. H.; Hwang, C. Y., et al., Antimicrobial Effects of Silver Nanoparticles (Vol 1, Pg 95, 2007). Nanomed.-Nanotechnol. Biol. Med. 2014, 10, 1119–1119.

25. Kandile, N. G.; Zaky, H. T.; Mohamed, M. I.; Mohamed, H. M., Silver Nanoparticles Effect on Antimicrobial and Antifungal Activity of New Heterocycles. Bull. Korean Chem. Soc. 2010, 31, 3530–3538.

26. Pal, S.; Tak, Y. K.; Song, J. M., Does the Antibacterial Activity of Silver Nanoparticles Depend on the Shape of the Nanoparticle? A Study of the Gram-Negative Bacterium Escherichia Coli. Applied and Environmental Microbiology 2007, 73, 1712–1720.

27. Raffi, M.; Mehrwan, S.; Bhatti, T. M.; Akhter, J. I.; Hameed, A.; Yawar, W.; ul Hasan, M. M., Investigations into the Antibacterial Behavior of Copper Nanoparticles against Escherichia Coli. Annals of Microbiology 2010, 60, 75–80.

28. Drelich, J.; Li, B. W.; Bowen, P.; Hwang, J. Y.; Mills, O.; Hoffman, D., Vermiculite Decorated with Copper Nanoparticles: Novel Antibacterial Hybrid Material. Applied Surface Science 2011, 257, 9435–9443.

29. AshaRani, P. V.; Mun, G. L. K.; Hande, M. P.; Valiyaveettil, S., Cytotoxicity and Genotoxicity of Silver Nanoparticles in Human Cells. ACS Nano 2009, 3, 279–290.

30. Kaegi, R.; Sinnet, B.; Zuleeg, S.; Hagendorfer, H.; Mueller, E.; Vonbank, R.; Boller, M.; Burkhardt, M., Release of Silver Nanoparticles from Outdoor Facades. Environmental Pollution 2010, 158, 2900–2905.

31. Chen, Z.; Meng, H. A.; Xing, G. M.; Chen, C. Y.; Zhao, Y. L.; Jia, G. A.; Wang, T. C.; Yuan, H.; Ye, C.; Zhao, F., et al., Acute Toxicological Effects of Copper Nanoparticles in Vivo. Toxicology Letters 2006, 163, 109–120.

32. Baek, Y. W.; An, Y. J., Microbial Toxicity of Metal Oxide Nanoparticles (Cuo, Nio, Zno, and Sb2o3) to Escherichia Coli, Bacillus Subtilis, and Streptococcus Aureus. Science of the Total Environment 2011, 409, 1603–1608.

33. Rosi, N. L.; Mirkin, C. A., Nanostructures in Biodiagnostics. Chemical Reviews 2005, 105, 1547–1562.

34. Stoimenov, P. K.; Klinger, R. L.; Marchin, G. L.; Klabunde, K. J., Metal Oxide Nanoparticles as Bactericidal Agents. Langmuir 2002, 18, 6679–6686.

35. Sun, C. Y.; Zhao, D.; Chen, C. C.; Ma, W. H.; Zhao, J. C., Tio2-Mediated Photocatalytic Debromination of Decabromodiphenyl Ether: Kinetics and Intermediates. Environ. Sci. Technol. 2009, 43, 157–162.

36. Kumar, B.; Smita, K.; Cumbal, L.; Debut, A.; Galeas, S.; Guerrero, V. H., Phytosynthesis and Photocatalytic Activity of Magnetite (Fe3o4) Nanoparticles Using the Andean Blackberry Leaf. Mater. Chem. Phys. 2016, 179, 310–315.

37. Comparelli, R.; Fanizza, E.; Curri, M. L.; Cozzoli, P. D.; Mascolo, G.; Passino, R.; Agostiano, A., Photocatalytic Degradation of Azo Dyes by Organic-Capped Anatase Tio2 Nanocrystals Immobilized onto Substrates. Applied Catalysis B-Environmental 2005, 55, 81–91.

38. Haick, H.; Paz, Y., Long-Range Effects of Noble Metals on the Photocatalytic Properties of Titanium Dioxide. Journal of Physical Chemistry B 2003, 107, 2319–2326.

39. Storz, G.; Imlay, J. A., Oxidative Stress. Current Opinion in Microbiology 1999, 2, 188–194.

40. Maisch, T.; Bosl, C.; Szeimies, R. M.; Lehn, N.; Abels, C., Photodynamic Effects of Novel Xf Porphyrin Derivatives on Prokaryotic and Eukaryotic Cells. Antimicrob. Agents Chemother. 2005, 49, 1542–1552.

41. Banerjee, M.; Mallick, S.; Paul, A.; Chattopadhyay, A.; Ghosh, S. S., Heightened Reactive Oxygen Species Generation in the Antimicrobial Activity of a Three Component Iodinated Chitosan-Silver Nanoparticle Composite. Langmuir 2010, 26, 5901–5908.

42. Raghupathi, K. R.; Koodali, R. T.; Manna, A. C., Size-Dependent Bacterial Growth Inhibition and Mechanism of Antibacterial Activity of Zinc Oxide Nanoparticles. Langmuir 2011, 27, 4020–4028.

43. Brayner, R.; Ferrari-Iliou, R.; Brivois, N.; Djediat, S.; Benedetti, M. F.; Fievet, F., Toxicological Impact Studies Based on Escherichia Coli Bacteria in Ultrafine Zno Nanoparticles Colloidal Medium. Nano Letters 2006, 6, 866–870.

44. Jones, N.; Ray, B.; Ranjit, K. T.; Manna, A. C., Antibacterial Activity of Zno Nanoparticle Suspensions on a Broad Spectrum of Microorganisms. Fems Microbiology Letters 2008, 279, 71–76.

45. Jalal, R.; Goharshadi, E. K.; Abareshi, M.; Moosavi, M.; Yousefi, A.; Nancarrow, P., Zno Nanofluids: Green Synthesis, Characterization, and Antibacterial Activity. Mater. Chem. Phys. 2010, 121, 198–201.

46. Padmavathy, N.; Vijayaraghavan, R., Enhanced Bioactivity of Zno Nanoparticles-an Antimicrobial Study. Science and Technology of Advanced Materials 2008, 9.

47. Chen, R.; Huo, L. L.; Shi, X. F.; Bai, R.; Zhang, Z. J.; Zhao, Y. L.; Chang, Y. Z.; Chen, C. Y., Endoplasmic Reticulum Stress Induced by Zinc Oxide Nanoparticles Is an Earlier Biomarker for Nanotoxicological Evaluation. ACS Nano 2014, 8, 2562–2574.

48. Yang, X.; Shao, H. L.; Liu, W. R.; Gu, W. Z.; Shu, X. L.; Mo, Y. Q.; Chen, X. J.; Zhang, Q. W.; Jiang, M. Z., Endoplasmic Reticulum Stress and Oxidative Stress Are Involved in Zno Nanoparticle-Induced Hepatotoxicity. Toxicology Letters 2015, 234, 40–49.

49. Wen, L.; Chen, L.; Zheng, S. M.; Zeng, J. F.; Duan, G. X.; Wang, Y.; Wang, G. L.; Chai, Z. F.; Li, Z.; Gao, M. Y., Ultrasmall Biocompatible Wo3-X Nanodots for Multi-Modality Imaging and Combined Therapy of Cancers. Advanced Materials 2016, 28, 5072–5079.

50. Adams, L. K.; Lyon, D. Y.; McIntosh, A.; Alvarez, P. J. J., Comparative Toxicity of Nano-Scale Tio2, Sio2 and Zno Water Suspensions. Water Science and Technology 2006, 54, 327–334.

51. Akhavan, O.; Ghaderi, E., Toxicity of Graphene and Graphene Oxide Nanowalls against Bacteria. ACS Nano 2010, 4, 5731–5736.

52. Niu, A.; Han, Y. J.; Wu, J. A.; Yu, N.; Xu, Q., Synthesis of One-Dimensional Carbon Nanomaterials Wrapped by Silver Nanoparticles and Their Antibacterial Behavior. Journal of Physical Chemistry C 2010, 114, 12728–12735.

53. Zhang, L. L.; Jiang, Y. H.; Ding, Y. L.; Povey, M.; York, D., Investigation into the Antibacterial Behaviour of Suspensions of Zno Nanoparticles (Zno Nanofluids). Journal of Nanoparticle Research 2007, 9, 479–489.

54. Zhang, W.; Ye, C.; De Luna, P.; Zhou, R., Snatching the Ligand or Destroying the Structure: Disruption of Ww Domain by Phosphorene. The Journal of Physical Chemistry C 2016.

55. Gu, Z.; De Luna, P.; Yang, Z.; Zhou, R., Structural Influence of Proteins Upon Adsorption to Mos 2 Nanomaterials: Comparison of Mos 2 Force Field Parameters. Physical Chemistry Chemical Physics 2017.

56. Tu, Y.; Lv, M.; Xiu, P.; Huynh, T.; Zhang, M.; Castelli, M.; Liu, Z.; Huang, Q.; Fan, C.; Fang, H., et al., Destructive Extraction of Phospholipids from Escherichia Coli Membranes by Graphene Nanosheets. Nature Nanotechnology 2013, 8, 594–601.

57. He, W. W.; Kim, H. K.; Wamer, W. G.; Melka, D.; Callahan, J. H.; Yin, J. J., Photogenerated Charge Carriers and Reactive Oxygen Species in Zno/Au Hybrid Nanostructures with Enhanced Photocatalytic and Antibacterial Activity. J. Am. Chem. Soc. 2014, 136, 750–757.

58. He, W. W.; Wu, H. H.; Warner, W. G.; Kim, H. K.; Zheng, J. W.; Jia, H. M.; Zheng, Z.; Yin, J. J., Unraveling the Enhanced Photocatalytic Activity and Phototoxicity of Zno/Metal Hybrid Nanostructures from Generation of Reactive Oxygen Species and Charge Carriers. Acs Applied Materials & Interfaces 2014, 6, 15527–15535.

59. Liang, Y.; Chen, Q.; Liu, Q.; Zhang, W.; Ding, R., Exogenous Silicon (Si) Increases Antioxidant Enzyme Activity and Reduces Lipid Peroxidation in Roots of Salt-Stressed Barley (Hordeum Vulgare L.). Journal of Plant Physiology 2003, 160, 1157–1164.

60. Fu, J.; Huang, B., Involvement of Antioxidants and Lipid Peroxidation in the Adaptation of Two Cool-Season Grasses to Localized Drought Stress. Environmental & Experimental Botany 2001, 45, 105–114.

61. Monopoli, M. P.; Aberg, C.; Salvati, A.; Dawson, K. A., Biomolecular Coronas Provide the Biological Identity of Nanosized Materials. Nature Nanotechnology 2012, 7, 779–786.

62. Duan, G. X.; Kang, S. G.; Tian, X.; Garate, J. A.; Zhao, L.; Ge, C. C.; Zhou, R. H., Protein Corona Mitigates the Cytotoxicity of Graphene Oxide by Reducing Its Physical Interaction with Cell Membrane. Nanoscale 2015, 7, 15214–15224.

63. Chong, Y.; Ge, C. C.; Yang, Z. X.; Garate, J. A.; Gu, Z. L.; Weber, J. K.; Liu, J. J.; Zhou, R. H., Reduced Cytotoxicity of Graphene Nanosheets Mediated by Blood-Protein Coating. ACS Nano 2015, 9, 5713–5724.

64. Zhou, R.; Berne, B. J.; Germain, R., The Free Energy Landscape for B Hairpin Folding in Explicit Water. Proceedings of the National Academy of Sciences 2001, 98, 14931–14936.

65. Das, P.; Li, J.; Royyuru, A. K.; Zhou, R., Free Energy Simulations Reveal a Double Mutant Avian H5n1 Virus Hemagglutinin with Altered Receptor Binding Specificity. Journal of computational chemistry 2009, 30, 1654–1663.

66. Xia, Z.; Clark, P.; Huynh, T.; Loher, P.; Zhao, Y.; Chen, H.-W.; Rigoutsos, I.; Zhou, R., Molecular Dynamics Simulations of Ago Silencing Complexes Reveal a Large Repertoire of Admissible ‘Seed-Less’ Targets. Scientific reports 2012, 2, 569.

67. Xiu, P.; Yang, Z.; Zhou, B.; Das, P.; Fang, H.; Zhou, R., Urea-Induced Drying of Hydrophobic Nanotubes: Comparison of Different Urea Models. The Journal of Physical Chemistry B 2011, 115, 2988–2994.

68. Li, J.; Liu, T.; Li, X.; Ye, L.; Chen, H.; Fang, H.; Wu, Z.; Zhou, R., Hydration and Dewetting near Graphite-Ch(3) and Graphite-Cooh Plates. J. Phys. Chem. B 2005, 109, 13639–48.

69. Zhou, R., Exploring the Protein Folding Free Energy Landscape: Coupling Replica Exchange Method with P3me/Respa Algorithm. J. Mol. Graph. Model. 2004, 22, 451–63.

70. Zhou, R.; Gao, H., Cytotoxicity of Graphene: Recent Advances and Future Perspective. Wiley Interdiscip Rev Nanomed Nanobiotechnol 2014, 6, 452–74.

71. Fitch, B. G.; Rayshubskiy, A.; Eleftheriou, M.; J., C. T.; Giampaga, M.; Zhestkov, Y.; Pitman, M. C.; Suits, F.; Grossfield, A.; Pitera, J., et al., Blue Matter: Strong Scaling of Molecular Dynamics on Blue Gene/L. Springe rBerlin Heidelberg: 2006.

72. Li, X.; Li, J.; Eleftheriou, M.; Zhou, R., Hydration and Dewetting near Fluorinated Superhydrophobic Plates. J. Am. Chem. Soc. 2006, 128, 12439–47.

73. Zhu, L.; Sheng, D.; Xu, C.; Dai, X.; Silver, M. A.; Li, J.; Li, P.; Wang, Y.; Wang, Y.; Chen, L., et al., Identifying the Recognition Site for Selective Trapping of (99)Tco4(-) in a Hydrolytically Stable and Radiation Resistant Cationic Metal-Organic Framework. J. Am. Chem. Soc. 2017, 139, 14873–14876.

74. Das, P.; Zhou, R., Urea-Induced Drying of Carbon Nanotubes Suggests Existence of a Dry Globule-Like Transient State During Chemical Denaturation of Proteins. J. Phys. Chem. B 2010, 114, 5427–30.

75. Kaminski, G. A.; Friesner, R. A.; Zhou, R., A Computationally Inexpensive Modification of the Point Dipole Electrostatic Polarization Model for Molecular Simulations. J. Comput. Chem. 2003, 24, 267–76.

76. Stirnemann, G.; Kang, S. G.; Zhou, R.; Berne, B. J., How Force Unfolding Differs from Chemical Denaturation. Proc. Natl. Acad. Sci. U. S. A. 2014, 111, 3413–8.

