## Supplementary material for "Robust Antibacterial Activity of Tungsten Oxide (WO_3-X_) Nanodots"

**Force field parameters**

The parameters for WO_3-x_ were first taken from UFF and then optimized to reproduce experimental water contact angle for 0 degree and quamtum mechanmical interaction distance with water.

The partial charge on O was determined by Mulliken analysis of a W_8_O_36_H_24_ cluster (**Figure S1A**) and Lowdin analysis of a WO_3-x_ slab model (**Figure S1B**). For the cluster, the geometry was optimized by using PBE functional, 6-311+G(d) basis set for O and H and LANL2DZ basis set for W, and then Mulliken analysis was conducted at the same level of theory. The slab model was constructed by cutting the [0 1 0] surface of a 1x2x1 supercell of the monoclinic WO_3_ crystal structure and removing the surface oxygen atoms. The vacuum thickness was 3.0 nm. The spurious interaction between periodic images was removed by the effective screening medium method. A 2x2x1 k-point mesh was used to sample the Brillouin zone. The rVV10 density functional was chosen, which has been shown to accurately describe both short-range and long-range interactions. The cutoff energy for the wavefunction was set to 75 Ry. Gaussian smearning was used with a spreading value of 0.02 Ry. Gaussian 09 and Quantum ESPRESSO 5.0 were used for the cluster and slab calculations, respectively.

The W charge in the cluster model ranges from +0.55 to +1.50 with an average of +1.00. The W charge in the slab model ranges from +0.81 to +1.06 with an average of +0.98. The values from the two methods are consistent with each other, and translate to -0.33 charge on O by using the chemical formula WO_3_. Since the WO3-x materials are oxygen-deficient, a slightly larger charge -0.35 for O was used, while the W charge was adjusted according to the total charge of the material. For example, in the W_18_O_49_ nanosheet, the W charge was +0.854.

To calculate the interaction energy between the WO_3-x_ slab and water, an optimized water molecule was placed at varying distances from the slab model. Several different water orientations and positions were included to reprensent interactions with W and O atoms on WO_3-x_, which are shown in **Figure S2**. The energies of isolated water and WO_3-x_ were then subtracted from the total energy to give the interaction energy. The QM methods were same as those used for the slab model.

Two methods were used to calculate the water contact angle. The first is dry-wall method. The free energy of adhesion between a water slab and the solid surface was computed by the double-decoupling method, *i.e.* the solid was grown both in the presence and absence of the water slab by alchemical transition, and the difference between the two free energy changes is the adhesion free energy. The water contact angle was then calculated by the relationship

$$W_{\text{adh}}=\gamma\left( 1+\cos\theta_{\text{Y}} \right)$$

Where $W_{\text{adh}}$ is the adhesion free energy per area, $\gamma$ is the surface tension of water and $\theta_{\text{Y}}$ is the Young’s contact angle. When $W_{\text{adh}}\geq2\gamma$, $\theta_{\text{Y}}=0$. The system for the free energy calculation consisted of 1069 TIP3P water molecules and a WO_3-x_ slab model constructed by cutting the [0 1 0] surface of 4x2x4 supercell of moniclinic WO_3_ and then removing surface oxygen. There were 21 alchemical states in the simulations. Firstly the vdW interaction was increased from zero to full strength in 10 steps, and then the electrostatics interaction was gradually grown in another 10 steps.

The second method was direct simulation, in which a water droplet on the surface was equilibrated and then the contact angle was measured. One droplet size containing ~4000 water molecules was used, and the WO_3-x_ slab model was constructed from 22x2x21 supercell of moniclinic WO_3_. The dimension of the simulation box was 16.1x12.0x16.2 nm^3^. After 10 ns equilibration, another 20 ns simulation trajectory was collected to compute the radial density distribution of water as a function of the distance to the surface, *z*. The radius of each water slice, *r*, was determined by fitting the radial density distribution to a sigmoidal function and finding the position with half of the bulk density. The slope of the *r*-*z* curve was then used to determine the contact angle. This method was mainly used for comparison, so no extrapolation to macroscopic droplet size was attempted.

The optimized force field parameters are shown in **Table S1**. The calculated adhesion surface energy is 107.0 dynes/cm, slightly larger than twice the surface tension of the TIP3P model (52.3 dynes/cm), which corresponds to a 0 degree water contact angle. The water contact angle calculated by the direct droplet simulation is also 0 degree. The interaction energy between water and WO_3-x_ is shown in **Table S2**. Because of the chemisorption nature, force field cannot accurately reproduce the QM interaction energy curve. Therefore we focused on improving the equilibrium interaction distance.

**Table S1.** Force field parameters

| Atom type | Charge (e) | σ (Angstrom) | ε (kcal/mol) |
| --- | --- | --- | --- |
| O | -0.350 | 3.1181 | 0.1795 |
| W | +0.796 ~ +0.908 | 2.3342 | 0.0670 |

**Table S2.** Comparison of water WO_3-x_ interaction energy and equilibrium distance calculated by QM, UFF and force field in this work.

| Structure | *R*_0_ (Angstrom) | | | *E*_b_ (kcal/mol) | | |
| --- | --- | --- | --- | --- | --- | --- |
|  | QM | UFF | This work | QM | UFF | This work |
| 1 | 2.170 | 2.511 | 2.344 | -22.29 | -9.88 | -8.71 |
| 2 | 2.329 | 2.657 | 2.419 | -18.22 | -0.97 | -3.61 |
| 3 | 2.252 | 1.954 | 1.999 | -3.47 | 0.97 | -1.52 |

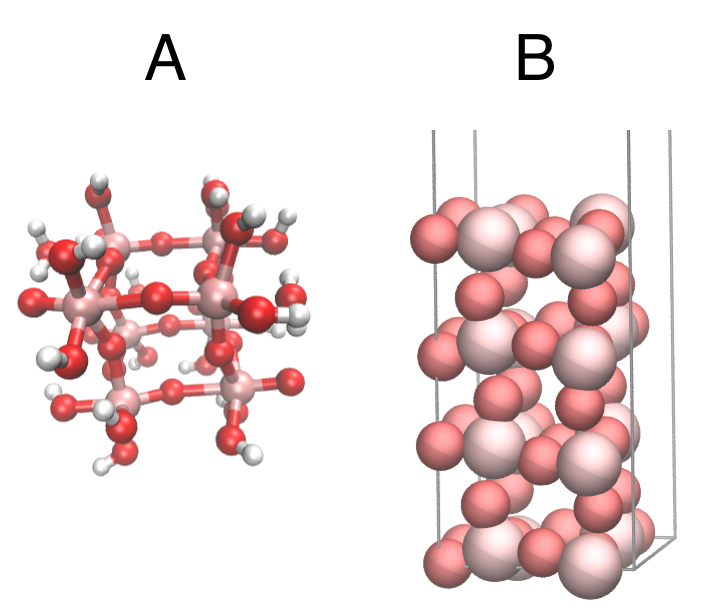

**Figure S1.** Structures used to determine the partial charges. (A) cluster model. (B) slab model. Pink, red, and white spheres are W, O, and H atoms, respectively.

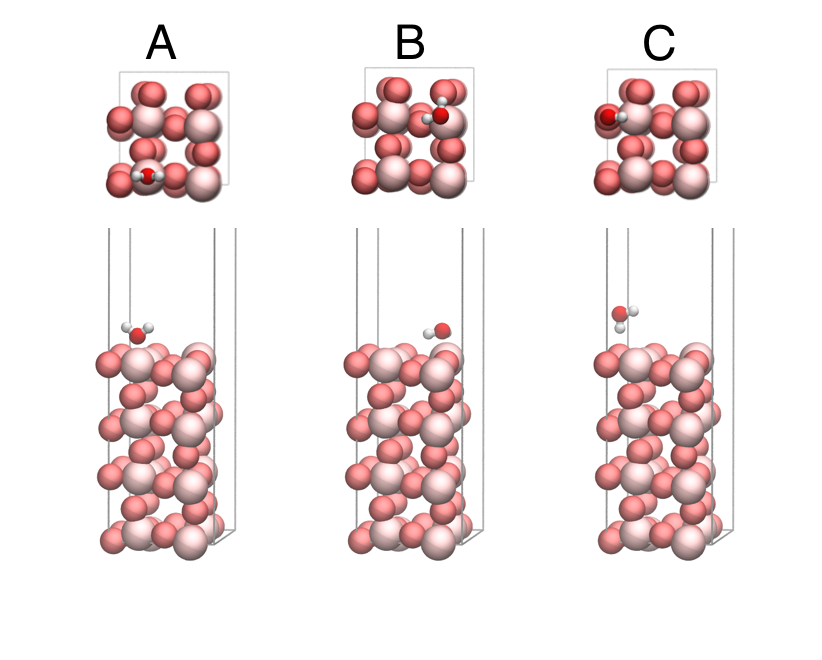

**Figure S2**. Front and top views of structures used to scan the interaction bewteen water and WO_3-x_. (A-C) are model 1, 2 and 3, respectively. Pink, red, and white spheres are W, O, and H atoms, respectively.

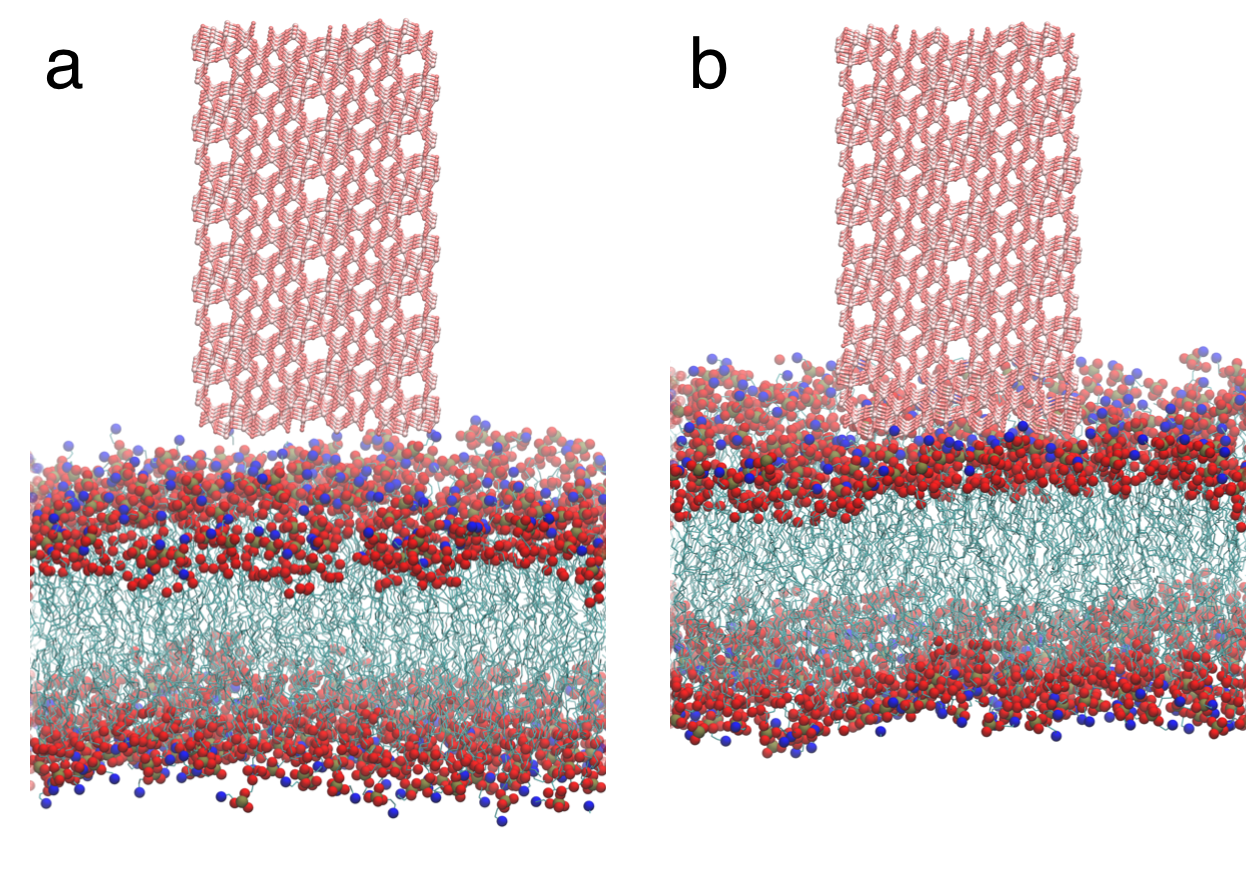

**Figure S3**. Initial (a) and final (b) conformations of the membrane-nanosheet simulations. The WO_3-x_ nanosheet attached to the membrane but cannot extract lipids out of the membrane.
